# Granzyme B prevents aberrant IL-17 production and intestinal pathogenicity in CD4^+^ T cells

**DOI:** 10.1101/2020.12.04.412056

**Authors:** Kristen L. Hoek, Michael J. Greer, Kathleen G. McClanahan, Ali Nazmi, M. Blanca Piazuelo, Kshipra Singh, Keith T. Wilson, Danyvid Olivares-Villagómez

## Abstract

CD4^+^ T cell activation and differentiation are important events that set the stage for proper immune responses. Many factors are involved in the activation and differentiation of T cells, and these events are tightly controlled to prevent unwanted and/or exacerbated immune responses that may harm the host. It has been well documented that granzyme B, a potent serine protease involved in cell-mediated cytotoxicity, is readily expressed by certain CD4^+^ T cells, such as regulatory T cells and CD4^+^CD8αα^+^ intestinal intraepithelial lymphocytes, both of which display cytotoxicity associated with granzyme B. However, because not all CD4^+^ T cells expressing granzyme B are cytotoxic, additional roles for this protease in CD4^+^ T cell biology remain unknown. Here, using a combination of *in vivo* and *in vitro* approaches, we report that granzyme B-deficient CD4^+^ T cells display increased IL-17 production. In the adoptive transfer model of intestinal inflammation, granzyme B-deficient CD4^+^ T cells triggered a more rapid disease onset than their WT counterparts, and presented a differential transcription profile. Similar results were also observed in granzyme B-deficient mice infected with *Citrobacter rodentium*. Our results suggest that granzyme B modulates CD4^+^ T cell differentiation, providing a new perspective into the biology of this enzyme.

## Introduction

Granzymes are a family of serine proteases expressed by many immune cell populations. There are five different granzymes in humans (A, B, H, K and M) and 11 in mice (A, B, C, D, E, F, G, K, L, M and N), which differ in function and substrate specificity.^1^ Most research has focused on granzymes A and B due to their hallmark role in CD8^+^ T and NK cell-mediated cytotoxicity against tumor cells or virus-infected cells.^2^ However, mounting evidence indicates that granzymes are also involved in other processes, including inflammatory responses,^3–5^ remodeling of the extracellular matrix,^6^ and cardiovascular disorders.^7^

Activated CD4^+^ T cells express granzyme B, and some of these cells exhibit lytic activity similar to CD8^+^ T cells.^8, 9^ Antigen-specific CD4^+^ T cell lytic activity mediated by granzyme B can be triggered by viral and bacterial infections.^10–12^ Similarly, some tumor-infiltrating CD4^+^ T cells also present anti-tumor cytotoxicity mediated by granzyme B.^13^ In addition, one of the mechanisms by which regulatory CD4^+^ T cells in humans and mice control immune responses is through granzyme B-mediated cell death.^14–18^

After activation, CD4^+^ T cells can differentiate into a diverse set of T helper (Th) cells depending on environmental cues and transcription factors, which ultimately leads Th cells to produce a specific cytokine profile.^19^ In certain populations of differentiated CD4^+^ T cells, granzyme B controls CD4^+^ T cell immune responses by initiating activation-induced cell death.^20^ Downregulation of granzyme B, for example when Th2 cells are cultured in the presence of vasoactive intestinal peptide, reduces Th2 cell death.^20^ Moreover, granzyme B expression in Th2 cells is enhanced by IL-10R signaling, which results in cell death.^21^ These studies clearly indicate that granzyme B is involved in controlling some immune CD4^+^ T cell responses. However, it is unclear whether this enzyme is actively involved during the process of Th differentiation.

In this report, we present data indicating that granzyme B is expressed during CD4^+^ T cell differentiation. Cells lacking granzyme B show a different cytokine profile than granzyme B-competent cells, primarily presenting increased IL-17 production. Moreover, using the adoptive transfer model of intestinal inflammation, we show that granzyme B-deficient CD4^+^ T cells activated *in vivo* have a different gene expression profile and display increased pathogenicity relative to WT cells. Similarly, we present evidence indicating that granzyme B deficiency predisposes mice to higher susceptibility to *Citrobacter rodentium* infection. These results indicate that granzyme B is required for proper CD4^+^ T cell differentiation, and in its absence, CD4^+^ T cells may acquire an aberrant phenotype characterized by increased IL-17 production and pathogenicity.

## Results

### Th0 and Th1, but not Th17 CD4^+^ T cells express granzyme B during differentiation

To determine the role of granzyme B in T cell activation and differentiation, we cultured naïve CD4^+^ T cells in Th0, Th1 and Th17 differentiation conditions. One and three days after culture, RNA was extracted to determine granzyme B expression relative to naïve CD4^+^ T cells. Under Th0 conditions, expression of granzyme B at day 1 displayed a slight increase above the levels observed for naïve CD4^+^ T cells (Figure 1a, top), but presented an almost three-fold increase at day 3 (Figure 1a, bottom). Th1 cells expressed a 3-fold increase in granzyme B mRNA over naïve T cells at day 1, which was maintained until day 3. On the other hand, CD4^+^ T cells under Th17 differentiation conditions did not induce granzyme B mRNA expression at either time point (Figure 1a).

**Figure 1.**
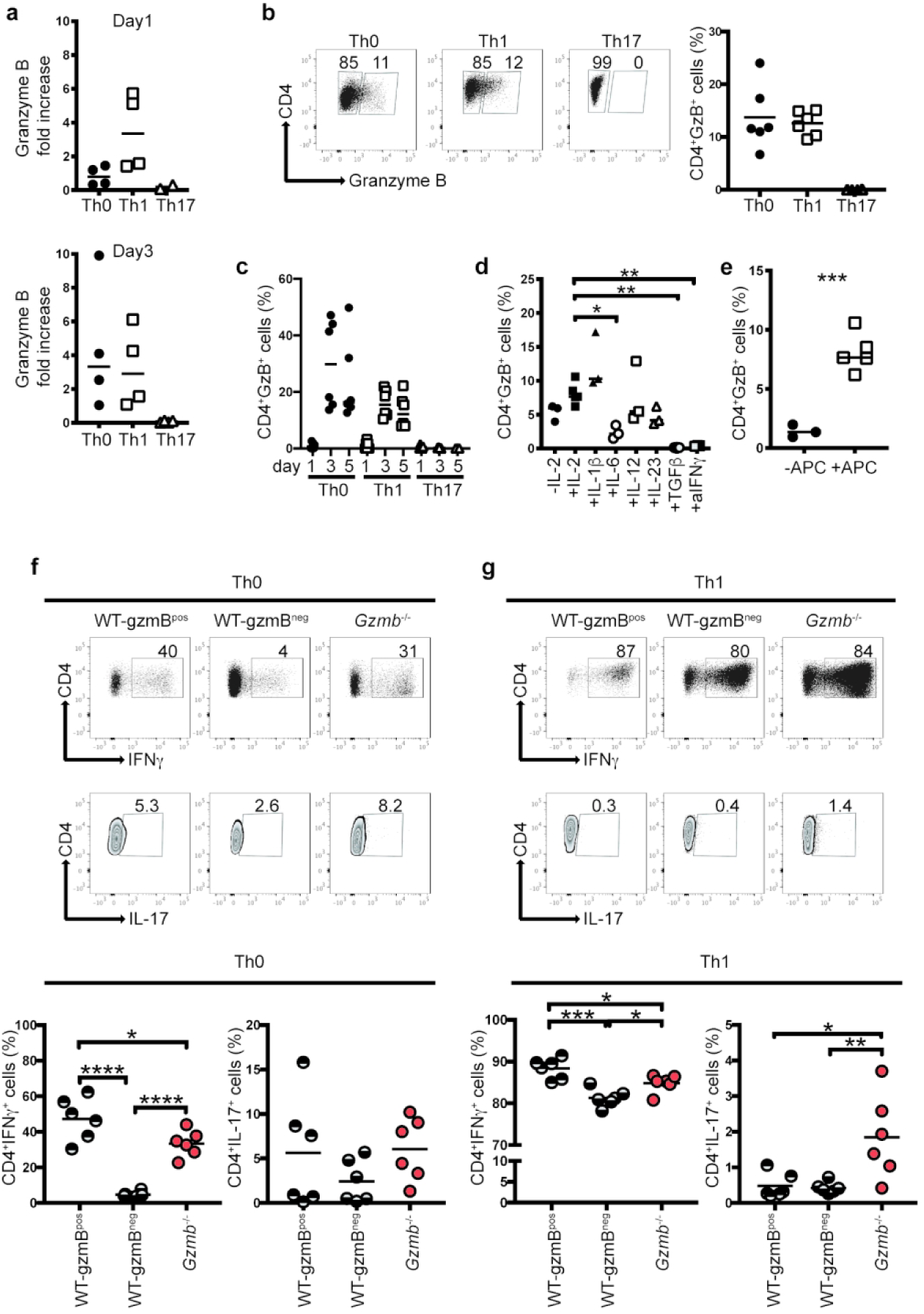
Granzyme B expression during T cell differentiation. (a) Naïve CD4^+^ T cells from WT mice were cultured in the presence of plate-bound anti-CD3 and soluble anti-CD28 under the indicated differentiation conditions, and in the absence of irradiated splenocytes. At day 1 (top) and 3 (bottom), cells were recovered, RNA extracted and granzyme B mRNA expression determined. Naïve CD4^+^ T cells without stimulation were used as reference. (b) Intracellular granzyme B in cells cultured as in (a), but in the presence of irradiated splenocytes. Left, representative dot plot (see Supplemental Figure 1a for gating strategy). Right, data summary. (c) Granzyme B expression in cells cultured in Th0, Th1 and Th17 conditions in the presence of irradiated splenocytes at different time points. (d) Cells cultured in Th0 conditions in the presence of irradiated splenocytes and the indicated individual cytokines, harvested 5 days after culture. (e) Cells cultured with IL-2, with or without irradiated splenocytes (APC). (f) and (g), intracellular IFNγ and IL-17 staining of WT and *Gzmb*^-/-^ CD4^+^ T cells cultured in Th0 or Th1 conditions, respectively, in the presence of irradiated splenocytes. Top, intracellular IFNγ and IL-17 staining of representative concatenated plots; bottom, data summary. Each dot represents an individual sample. Data are representative of at least 3 independent experiments, performed in triplicate. n=6. For (d), (f) and (g), *P<0.05; **P<0.01; ***P<0.001; ****P<0.001, One-way ANOVA; For (e), ***P<0.001, Student’s t test.

Around 13% of Th0 and Th1 cells expressed granzyme B intracellularly at day 5 (Figure 1b; see Supplemental Figure 1a for gating strategy), whereas this enzyme was not detected in Th17 cells, confirming the results for granzyme B mRNA expression. Although granzyme B mRNA was detected at day 1 in Th1 cells, granzyme B protein was not, but the protein was conspicuously expressed at day 3 and day 5 (Figure 1c). These results suggest a lag between granzyme B mRNA transcription and translation. A similar granzyme B protein expression profile was detected for Th0 cells (Figure 1c). On the other hand, Th17 cells did not express granzyme B at any of the time points analyzed (Figure 1c). Similarly, Th2 cells did not express granzyme B (Supplemental Figure 2b).

Because Th1 and Th17 cultures contain different cytokine/antibody cocktails, and granzyme B is expressed in the former, but not in the latter, we investigated what factors present in these culture conditions induce or repress granzyme B expression. For this purpose, we activated CD4^+^ T cells under Th0 conditions (anti-CD3/CD28, IL-2, and irradiated spleen cells), in the presence of individual factors included in Th1 and Th17 differentiation cocktails (Figure 1d). IL-2 alone induced ∼7.5% of CD4^+^ T cells to express granzyme B, which was not significantly different from cells activated without IL-2 (background levels), or in the presence of IL-12, IL-1β, and IL-23, the last two of which are involved in Th17 differentiation. However, the Th17-inducing cytokines IL-6 and TGFβ, as well as the blocking anti-IFNγ antibody, significantly ablated granzyme B expression. These results indicate that CD4^+^ T cell activation induces granzyme B activation, whereas some cytokines and anti-IFNγ inhibit its expression.

Because most of our differentiation cultures included spleen cells as a source of APC, we determined whether these cells influence granzyme B expression during CD4^+^ T cell activation. As shown in Figure 1e, CD4^+^ T cell activation in the absence of APC (but including IL-2), resulted in less than 2% of CD4^+^ T cells expressing granzyme B, whereas irradiated spleen APC significantly increased granzyme B expression.

Overall, these results indicate that granzyme B is expressed during Th0 and Th1 differentiation, with irradiated APC being an important expression-enhancing factor. On the other hand, Th17 and Th2 differentiation conditions ablate granzyme B expression. This effect can be attributed to IL-6, TGFβ, and anti-IFNγ.

As shown in Figure 1b, during Th0 and Th1 differentiation, two populations of CD4^+^ T cells can be distinguished: WT-gzmB^pos^ and WT-gzmB^neg^. To determine whether these populations present different cytokine profiles, we determined the frequencies of IFNγ- and IL- 17-producing cells in each of these subpopulations. Around half of Th0 WT-gzmB^pos^ expressed IFNγ, whereas only around 5% of the WT-gzmB^neg^ CD4^+^ T cells were positive for IFNγ (Figure 1f). Th0 cells derived from granzyme B-deficient (*Gzmb*^-/-^) mice presented decreased levels of IFNγ-producing cells in comparison to WT-gzmB^pos^ cells, but significantly more IFNγ^+^ cells than WT-gzmB^neg^ cells (Figure 1f). On the other hand, IL-17 production was not significantly different among Th0 cells from any of the CD4^+^ T cell populations.

Th1 differentiation induced high levels of IFNγ^+^ cells in all CD4^+^ T cell populations analyzed; however, WT-gzmB^pos^ cells presented the highest levels of IFNγ^+^ cells in comparison to WT-gzmB^neg^ or cells derived from *Gzmb*^-/-^ mice (Figure 1g). Interestingly, under Th1 conditions, cells from *Gzmb*^-/-^ mice presented significantly greater levels of IL-17^+^ cells when compared to WT-gzmB^pos^ and WT-gzmB^neg^ cells (Figure 1g). These results suggest that granzyme B may be involved in preventing IL-17 production when CD4^+^ T cells are cultured under Th1 conditions.

CD4^+^ T cells cultured in Th2 differentiation conditions, which contains anti-IFNγ, lack granzyme B expression (Supplemental Figure 1b); therefore, as expected, promiscuous cytokine expression in granzyme B-deficient CD4^+^ T cells was not observed in Th2 differentiation conditions (Supplemental Figure 1c).

### *In vivo* activation of granzyme B-deficient CD4^+^ T cells results in increased pathogenicity

To further understand the role of granzyme B in CD4^+^ T cell differentiation, we adoptively transferred naïve CD4^+^CD45RB^hi^ T cells from WT or *Gzmb*^-/-^ mice into *Rag2^-/-^* recipient mice. Because it is well-established that transfer of naïve CD4^+^ T cells into immunocompromised recipients results in chronic intestinal inflammation, ^22, 23^ this set up allows us to: 1) determine whether granzyme B-deficiency alters CD4^+^ T cell pathogenicity, and 2) whether absence of granzyme B influences the cytokine profile of CD4^+^ T cells activated *in vivo* (discussed in the next section).

In our animal colony, transfer of naïve CD4^+^ T cells from WT mice typically results in disease symptoms by 6 weeks after transfer. However, when naïve CD4^+^ T cells from *Gzmb*^-/-^ mice were transferred into *Rag2*^-/-^ mice, weight loss was evident starting at 14 days after transfer, and by day 21 these mice had lost around 20% of their original weight (Figure 2a). At this point, the experiments were terminated to comply with our institution’s animal welfare regulations. In contrast, at day 21, *Rag2*^-/-^ mice that received WT cells were either gaining or maintaining their weight (Figure 2a). Weight loss was accompanied by increased disease severity characterized by diarrhea, pilo-erection, and hunching (Figure 2b); increased intestinal pathology based on immune cell infiltration, loss of goblet cells, epithelial damage and hyperplasia (Figure 2c); and donor-derived CD4^+^ T cell reconstitution (Figure 2d). To confirm that the pathogenicity observed in granzyme B-deficient CD4^+^ T cells is an intrinsic effect, we adoptively transferred naïve CD4^+^ T cells from littermate *Gzmb*^+/-^ and *Gzmb*^-/-^ mice into *Rag2*^-/-^ recipient mice. Although disease development was delayed for almost a week, *Gzmb*^-/-^ mice displayed earlier disease onset and increased colon pathology compared to *Gzmb*^+/-^ littermate controls (Supplemental Figure 2a), indicating that the pathogenicity observed is intrinsic to CD4^+^ T cells.

**Figure 2.**
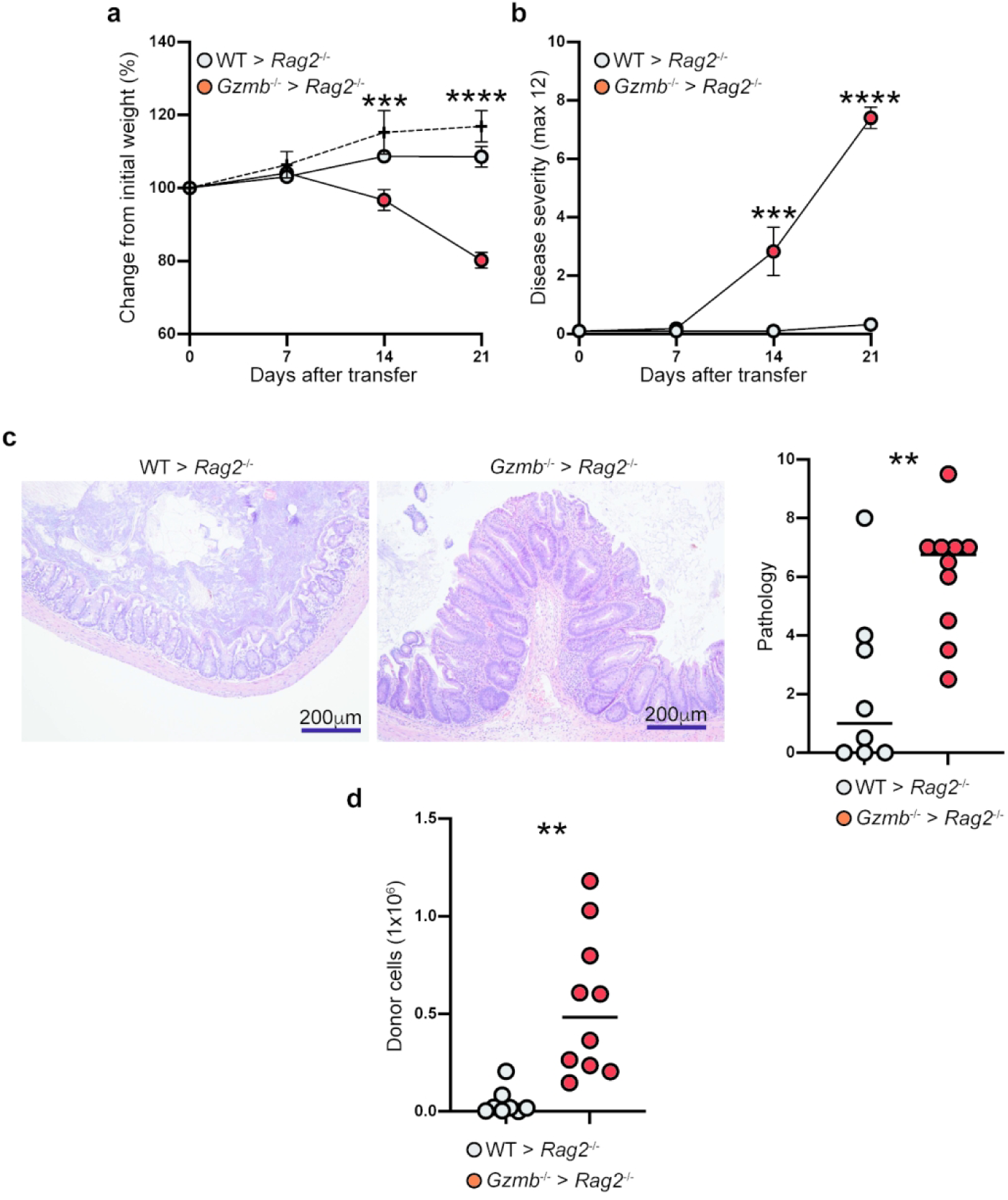
*In vivo* activated granzyme B-deficient CD4^+^ T cells display increased pathogenicity. 1×10^5^ CD4^+^CD45RB^hi^ T cells from WT or *Gzmb*^-/-^ donor mice were adoptively transferred into *Rag2*^-/-^ recipient mice. (a) Mice were weighed weekly, and (b) monitored for signs of disease. Twenty-one days after transfer, (c) colon pathology was scored (left, representative micrographs; right, data summary), and (d) donor-cell reconstitution of the IEL compartment in the colon was determined. For (a), dotted lines represent untreated *Rag2*^-/-^ mice. Each dot represents an individual mouse. Data are representative of at least 3 independent experiments. n=8-10. **P<0.01; ***P<0.001; ****P<0.0001; Student’s t-test.

To investigate whether the recipient mice’s granzyme B status is important for disease development, we adoptively transferred naïve CD4^+^ T cells from WT or *Gzmb*^-/-^ donor mice into *Rag2*^-/-^ *Gzmb*^-/-^ recipient mice. Donor CD4^+^ T cells from WT mice did not cause early weight loss in the double knock-out recipient mice, while CD4^+^ T cells deficient in granzyme B caused similar weight loss as in *Rag2*^-/-^ recipient mice (Supplemental Figure 2b and Figure 2a). Weight loss in mice receiving granzyme B-deficient CD4^+^ T cells was accompanied by increased disease severity (Supplemental Figure 2c), colon pathology (Supplemental Figure 3d), and donor-derived CD4^+^ T cell reconstitution (Supplemental Figure 3e). Therefore, the granzyme B status in the recipient mice is irrelevant for the intrinsic pathogenicity of granzyme B-deficient CD4^+^ T cells.

In summary, our data indicate that absence of granzyme B in CD4^+^ T cells activated *in vivo* results in accelerated pathogenicity and increased cell reconstitution.

### Granzyme B-deficient CD4^+^ T cells present increased IL-17 production *in vivo*

To determine whether granzyme B deficiency results in differential CD4^+^ T cell cytokine profiles *in vivo*, we adoptively transferred naïve CD4^+^CD45RB^hi^ T cells from WT or *Gzmb*^-/-^ mice into *Rag2*^-/-^ recipient mice. Three weeks after transfer, donor-derived cells were recovered from the MLN and lamina propria and analyzed for IFNγ and IL-17 expression. As shown in Figure 3, IFNγ was the predominant cytokine produced by donor CD4^+^ T cells derived from WT and *Gzmb*^-/-^ mice, with similar levels between WT and *Gzmb*^-/-^ CD4^+^ T cells (Figure 3a and 3b). However, CD4^+^ T cells from *Gzmb*^-/-^ donor mice presented significantly increased IL-17 production in comparison to WT CD4^+^ T cells (Figure 3a and 3b). It has been reported that pathogenesis in colitis requires the development of Th17 cells expressing IFNγ.^24^ As shown in Figure 3a and 3b, the frequency of IL-17^+^IFNγ^+^ donor-derived CD4^+^ T cells was similar among both groups in the MLN and lamina propria, suggesting that these cells may not be responsible for the accelerated colitis development observed. Albeit less pronounced, cytokine profiles were also similar when littermate *Gzmb*^+/-^ and *Gzmb*^-/-^ donor mice were used (Supplemental Figure 3).

**Figure 3.**
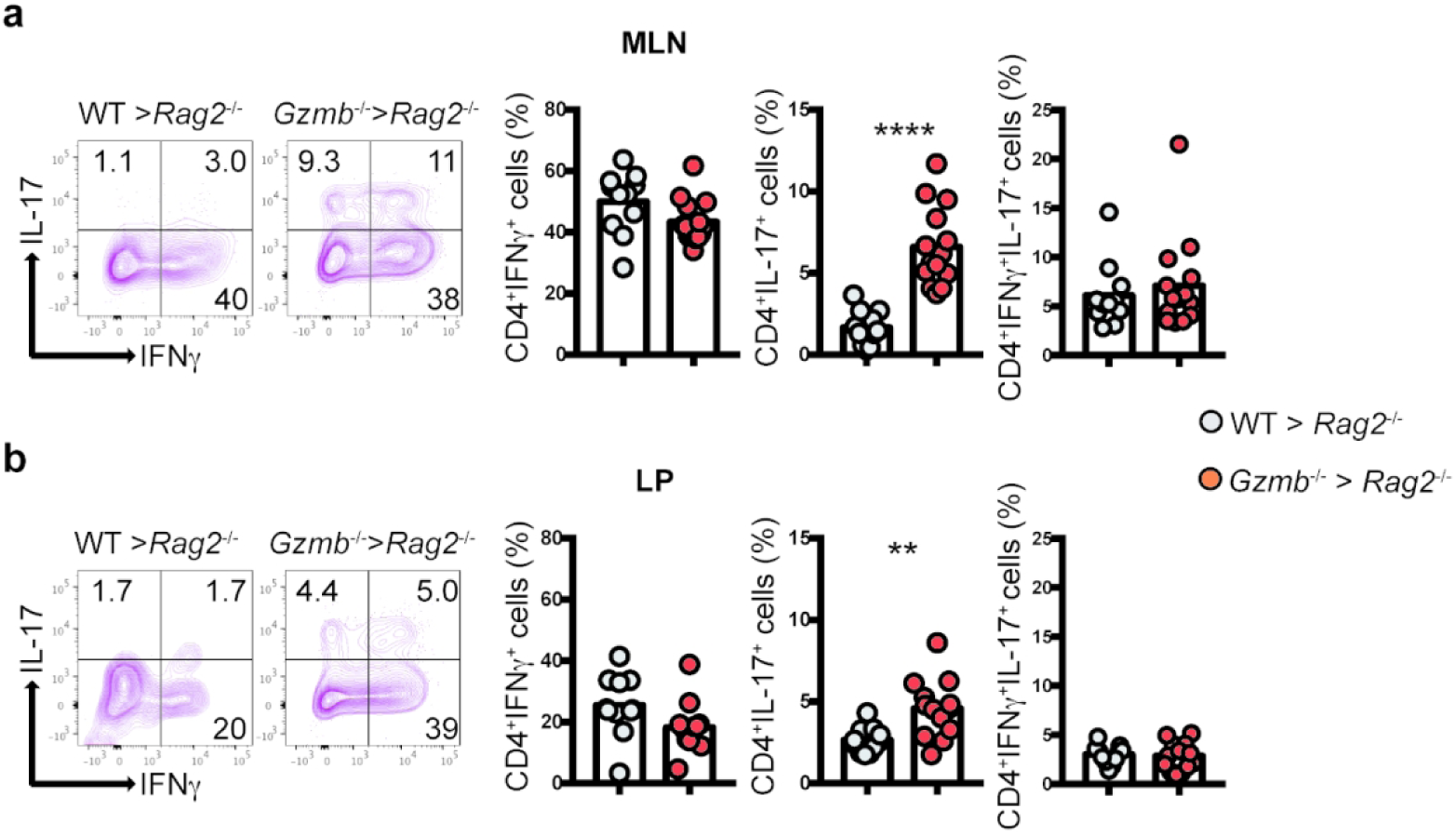
Granzyme B-deficient CD4^+^ T cells display skewed IL-17 differentiation *in vivo.* 1×10^5^ CD4^+^CD45RB^hi^ T cells from WT or *Gzmb*^-/-^ donor mice were adoptively transferred into *Rag2*^-/-^ recipient mice. Three weeks after transfer, donor-derived cells were recovered from (a) MLN, and (b) lamina propria, and their IFNγ/IL-17 profile was determined by intracellular staining. Dot plots indicate a representative sample. Live, TCRβ^+^CD4^+^ cells are displayed. Graphs represent the summary. Each dot indicates an individual mouse. For MLN, n=11-13; for lamina propria, n=7-11. Data are representative of at least 3 independent experiments. **P<0.01; ****P<0.001. Student’s t-Test.

In summary, *in vivo* activated CD4^+^ T cells deficient in granzyme B presented increased IL-17 production. These results suggest that absence of granzyme B results in skewed *in vivo* differentiation.

### Granzyme B deficient CD4^+^ T cells present a distinct gene expression profile

Because granzyme B-deficient CD4^+^ T cells had increased IL-17 production when activated *in vivo*, we investigated whether these cells presented a distinct gene expression profile in comparison to *in vivo* activated CD4^+^ T cells from WT mice. For this purpose, CD4^+^CD45RB^hi^ T cells from WT and *Gzmb*^-/-^ donor mice were adoptively transferred into *Rag2*^-/-^ mice. Three weeks after transfer, CD4^+^ T cells were recovered from the MLN, their RNA extracted and gene expression profile determined by RNAseq. As shown in Figure 4a, WT and granzyme B-deficient CD4^+^ T cells presented significantly different gene expression profiles. Examples of genes encoding transcription factors or molecules associated with gene expression upregulated in CD4^+^ T cells derived from *Gzmb*^-/-^ mice include Special AT-rich Binding protein (*Satb1*), a chromatin remodeling protein involved in T cell development and implicated in Th2 and regulatory T cell responses;^25^ Krüppel-like transcription factor 13 (*Klf13*), which has been implicated in T cell survival; ^26^ nuclear receptor subfamily 1, group D, member 1 (*Nr1d1*), which in human T cells is important for circadian cycles;^27^ and the Th17 master transcription factor *Rorc*. We also performed gene set enrichment analysis to determine whether WT and granzyme B-deficient CD4^+^ T cells recovered from recipient mice segregated in specific groups. As indicated in Figure 4b, GSEA indicated that these two CD4^+^ T cell populations presented distinct differential gene expression within several GSEA groups (Figure 4b). These results indicate that granzyme B-deficient CD4^+^ T cells, in the adoptive transfer system, follow a distinct differentiation pathway compared to WT CD4^+^ T cells.

**Figure 4.**
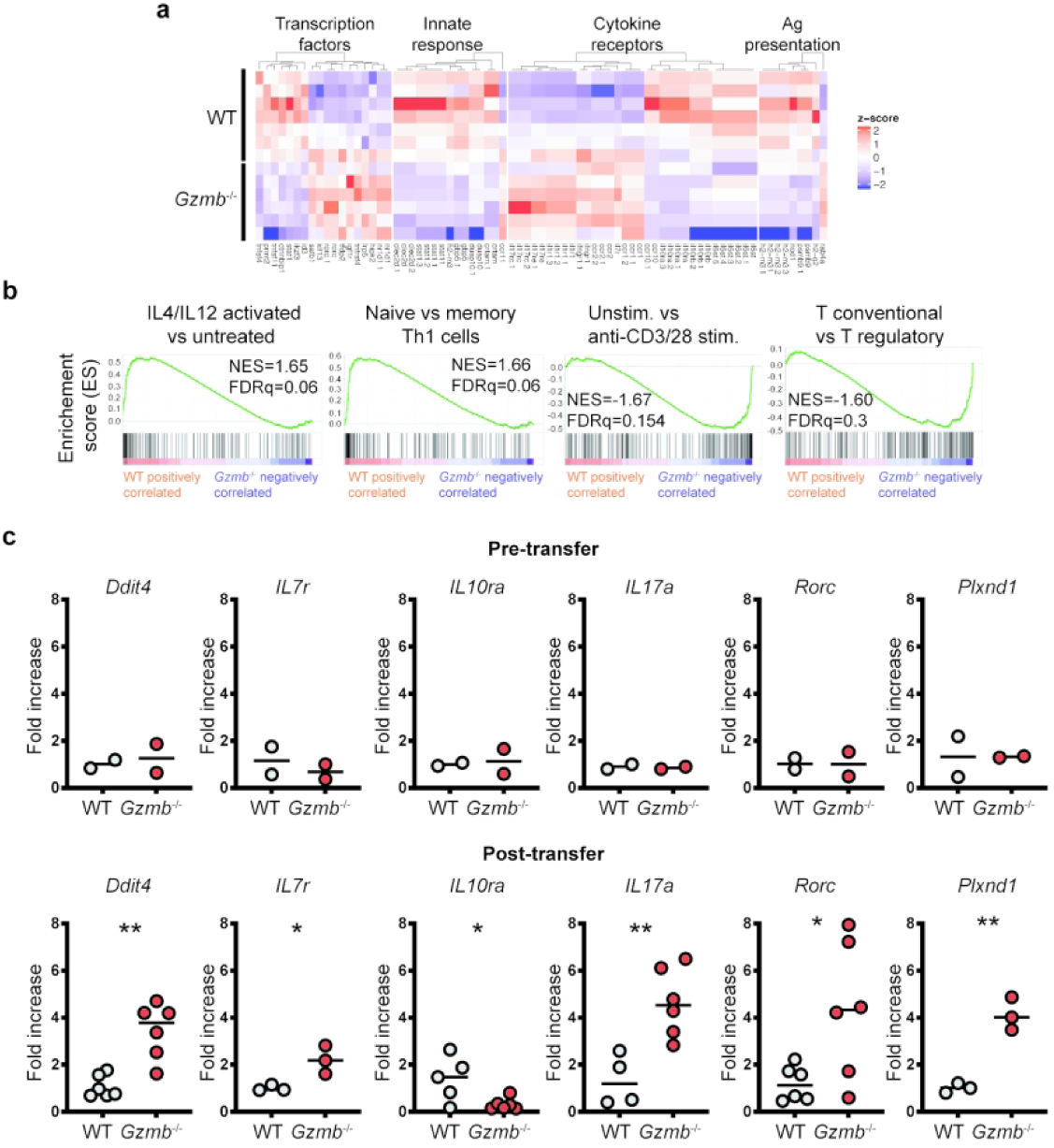
Granzyme B-deficient CD4^+^ T cells activated *in vivo* display a distinct gene expression profile. 1×10^5^ CD4^+^CD45RB^hi^ T cells from WT or *Gzmb*^-/-^ donor mice were adoptively transferred into *Rag2*^-/-^ recipient mice. Three weeks after transfer, donor-derived cells were recovered from the MLN, sorted for live TCRβ^+^CD4^+^, their RNA isolated and sequenced. (a) Representative heat map. (b) Representative gene set enrichment analysis. The green line on the plots indicate the enrichment score; the black lines show where the genes related to the pathway are located in the ranking; the legend at the bottom indicates whether the expression of the genes correlate more with CD4^+^ T cells derived from WT or *Gzmb*^-/-^ mice. (c) Real-time PCR validation of selected genes. For day 0, n=2; for day 21, n=3-6. *P<0.05; **P<0.01. Student’s t-Test.

We validated the results generated from the RNAseq analysis by comparing the expression of selected genes in naïve CD4^+^ T cells (Figure 4c, top), and donor-derived cells recovered 21d post-adoptive transfer (Figure 4c, bottom). Hallmark genes involved in Th17 responses, such as *Rorc* and *Il17a*, were similarly expressed in naïve CD4^+^ T cells, regardless of the granzyme B status, but their expression was significantly increased in recovered granzyme B-deficient CD4^+^ T cells. *Ddit4*, a gene important for optimal T cell proliferation,^28^ is highly expressed in T cells from multiple sclerosis patients, and is important in Th17 differentiation.^29^ *Ddit4* expression was 3-fold higher in activated CD4^+^ T cells derived from *Gzmb*^-/-^ mice in comparison to WT CD4^+^ T cells (Figure 4c). Granzyme B-deficient CD4^+^ T cells presented a 2-fold increase in *Il7r* expression (Figure 4c), which mediates IL-7 signaling, a well-known lymphopoietic cytokine also expressed in T cells undergoing homeostatic expansion.^30^ Plexin D1, encoded by the *Plxnd1* gene, binds semaphorins^31^ and is critical for directing thymocyte migration.^32^ Although the role of *Plxnd1* is not well known in mature T cell biology, granzyme B-deficient CD4^+^ T cells presented approximately a 4-fold increase in expression. Interestingly, donor CD4^+^ T cells derived from *Gzmb*^-/-^ mice presented decreased *Il10ra* expression in comparison to WT counterparts (Figure 4c).

In terms of IFNγ expression, the total count read average from the RNAseq analysis for this gene were 2940.7±1332 and 2609.3±1348.9 (P=0.66) for cells derived from WT and *Gzmb*^-/-^ CD4^+^ T cells, respectively, indicating that IFNγ expression is similar among the two groups. These results confirm what we observed with intracellular staining for this cytokine (Figure 3).

Overall, these results show that in the adoptive transfer model of CD4^+^ T cell activation, granzyme B deficiency results in a distinct gene expression profile, which is characterized by genes relevant to Th17 differentiation and T cell proliferation.

### Granzyme B-deficient CD4^+^ T cells possess better reconstitution capability

As indicated in Figure 2d, granzyme B-deficient CD4^+^ T cells showed greater reconstitution in the colon than WT CD4^+^ T cells when transferred into Rag-2^-/-^ recipient mice. However, this observation could be due to inflammation-driven proliferation, which was more prevalent in *Rag2*^-/-^ mice receiving CD4^+^ T cells from *Gzmb*^-/-^ mice. To further determine the reconstitution potential of CD4^+^ T cells derived from WT and *Gzmb*^-/-^ mice, we performed competitive adoptive T cell transfer experiments. For this purpose, naïve CD4^+^ T cells from WT CD45.1 and *Gzmb*^-/-^ CD45.2 mice (1×10^5^ each) were adoptively co-transferred into the same *Rag2*^-/-^ recipient mice. Of the total donor-derived cells obtained from the MLN, granzyme B-deficient CD4^+^ T cells presented ∼60% reconstitution, whereas WT-derived cells showed ∼40% reconstitution (Figure 5a). To determine whether increased cell reconstitution by granzyme B-deficient CD4^+^ T cells correlated with increased cell division, we stained the cells for Ki67 as a surrogate marker for cell division. As shown in Figure 5b, there was higher percentage of Ki67^+^ cells in granzyme-B deficient donor-derived cells. To further confirm these results, we cultured donor-derived cells without any stimulation, and measured their division potential by CFSE dilution after 3 days. CD4^+^ T cells from *Gzmb*^-/-^ mice showed a slight increase in cell division compared to cells derived from WT mice (Figure 5c).

**Figure 5.**
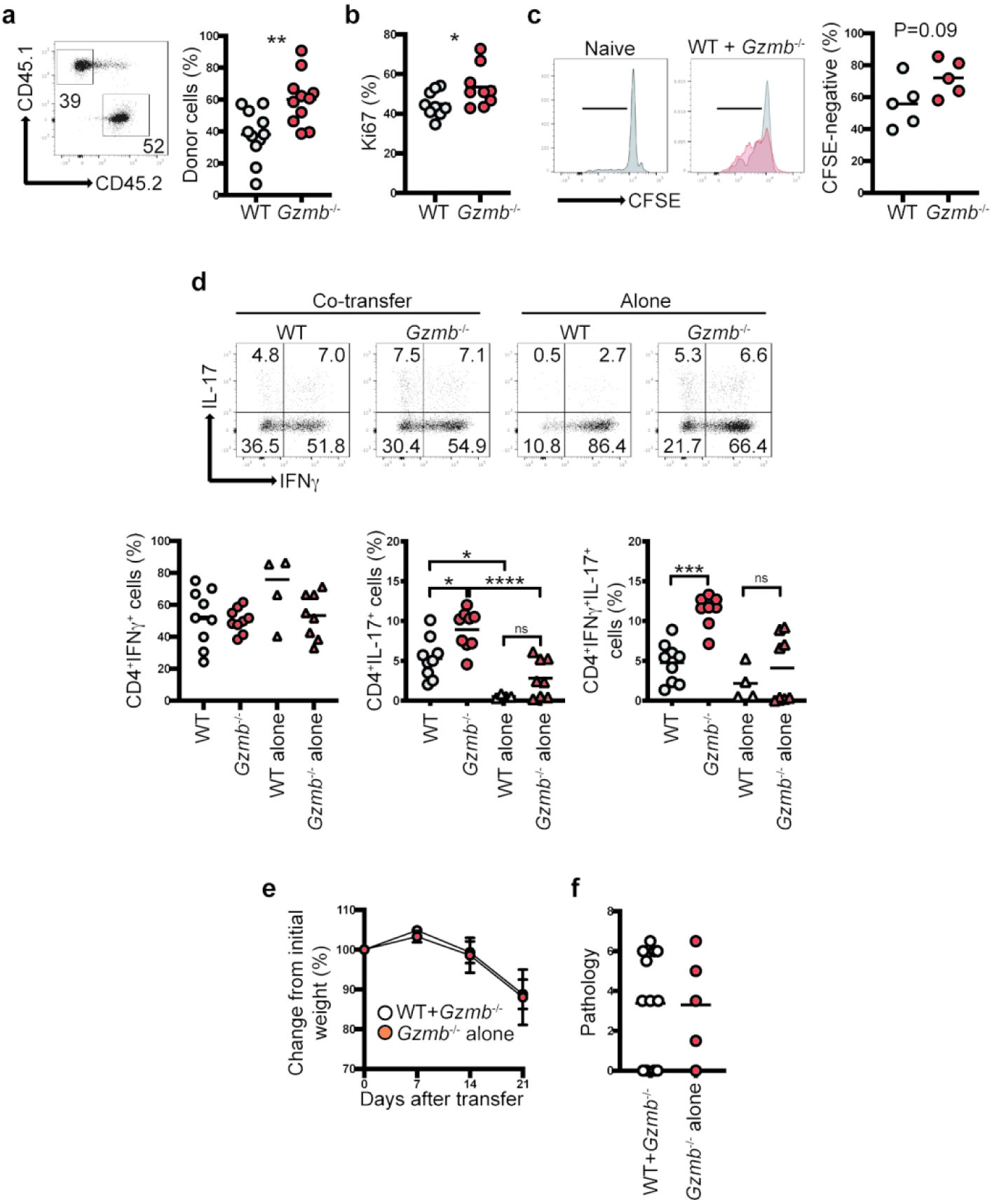
Granzyme B-deficient CD4^+^ T cells present greater *in vivo* reconstitution. 1×10^5^ CD4^+^CD45RB^hi^ T cells from WT (CD45.1) and *Gzmb*^-/-^ (CD45.2) donor mice were co-transferred into the same *Rag2*^-/-^ recipient mice. As controls, cells from the same donors were independently transferred into *Rag2*^-/-^ recipient mice. Three weeks after transfer, (a) MLN reconstitution (right, representative dot plot displaying live, TCRβ^+^CD4^+^ cells; left, summary), and (b) Ki67 staining were determined. (c) *In vitro* proliferation potential measured by CFSE dilution. Histograms indicate representative samples. Gray, WT; Red, *Gzmb*^-/-^. The gates indicate CFSE-negative cells; graph represents summary. (d) IFNγ/IL-17 production was determined from cells as in (a); graphs represent single IFNγ^+^ or IL-17^+^ (left and middle, respectively), and IFNγ^+^IL-17^+^ cells (right). (e) *Rag2*^-/-^ mice that received CD4^+^ T cells from both WT/*Gzmb*^-/-^ or only *Gzmb*^-/-^ mice were weighed weekly. (f) Colon pathology was determined 3 weeks post transfer. For (a) n=11; (b) n=9; (c) n=5; (d) n=4-9; (e), n=5-9. Each dot represents an individual mouse. Data are representative of at least 2 independent experiments. *P<0.05; **P<0.01; ***P<0.001; ****P<0.0001. For (a) and (b) Student’s t-Test; for (d) One-Way ANOVA.

We also analyzed the cytokine profile of the cells derived from the competitive adoptive CD4^+^ T cell approach. Although the IFNγ profile of co-transferred cells was similar to cells transferred alone, co-transferred WT cells presented increased IL-17 production in comparison to WT CD4^+^ T cells transferred alone (Figure 5d, left and middle graphs). These results raise the possibility that granzyme B-deficient CD4^+^ T cells influenced the behavior of granzyme B-competent CD4^+^ T cells. Interestingly, co-transferred granzyme B-deficient CD4^+^ T cells displayed a higher percentage of IFNγ^+^IL-17^+^ cells in comparison to WT cells (Figure 5d, right graph). *Rag2*^-/-^ mice co-transferred with WT and GzB^-/-^ CD4^+^ T cells displayed similar weight loss (Figure 5e) and colon pathology (Figure 5f) to mice transferred only with CD4^+^ T cells from *Gzmb* ^-/-^ mice, suggesting that the increased pathogenicity of the latter cells was maintained even in the presence of granzyme-competent CD4^+^ T cells.

### Granzyme B-deficient mice present normal cellularity and IFNγ /IL-17 production

Since *in vivo* activated granzyme B-deficient CD4^+^ T cells possess differential gene expression and cytokine profiles, it is possible that these differences are also observed in naïve cells. To investigate this possibility, we enumerated the CD4^+^ T cell cellularity of spleen, MLN, and IEL compartments from WT and *Gzmb*^-/-^ mice. As indicated in Supplemental Figure 4a, frequencies and numbers of TCRβ^+^CD4^+^ T cells were similar among the two groups of mice. *Ex vivo* IFNγ and IL-17 production by non-stimulated (ns) or stimulated (s) naïve CD4^+^ T cells derived from the MLN and IEL showed no distinguishable difference between cells from WT and *Gzmb*^-/-^ mice (Supplemental Figure 4b).

In summary, naïve WT and *Gzmb*^-/-^ animals appear to have similar CD4^+^ T cell cellularity and basal IFNγ and IL-17 production in peripheral lymphoid organs and the intestines.

### Granzyme B deficiency results in increased disease severity during *Citrobacter rodentium* infection

As demonstrated in the previous section, naïve *Gzmb*^-/-^ mice have a normal CD4^+^ T cell compartment compared to WT mice. However, because we observed that activated T cells from *Gzmb*^-/-^ mice became more pathogenic and had a distinct gene expression profile in the adoptive transfer model of colitis, we investigated whether *Gzmb*^-/-^ mice respond properly to antigenic stimulus. For this purpose, we infected WT and *Gzmb* ^-/-^ mice with the extracellular bacterium *Citrobacter rodentium*, which preferentially colonizes the colon of mice and induces an IL-17/IL-22-based immune response. After infection, WT mice maintained similar weight throughout the course of the experiment; however, starting at 9 days post-infection, *Gzmb*^-/-^ mice lost weight, reaching approximately ∼20% loss of the starting weight by day 12 (Figure 6a). *Gzmb*^-/-^ mice also presented other signs of disease, such as diarrhea, hunched posture, and pilo-erection, which were mostly absent in WT mice (Figure 6b). Despite similar colonic bacterial burden (Figure 6c), *Gzmb*^-/-^ mice developed greater pathology characterized by increased infiltration and epithelial injury than WT mice (Figure 6d). Analysis of cytokine production showed that in the spleen and LP of infected *Gzmb*^-/-^ mice, there was a significant increase in CD4^+^ T cells expressing IL-17, while IFNγ production was only increased in the MLN (Figure 6e).

**Figure 6.**
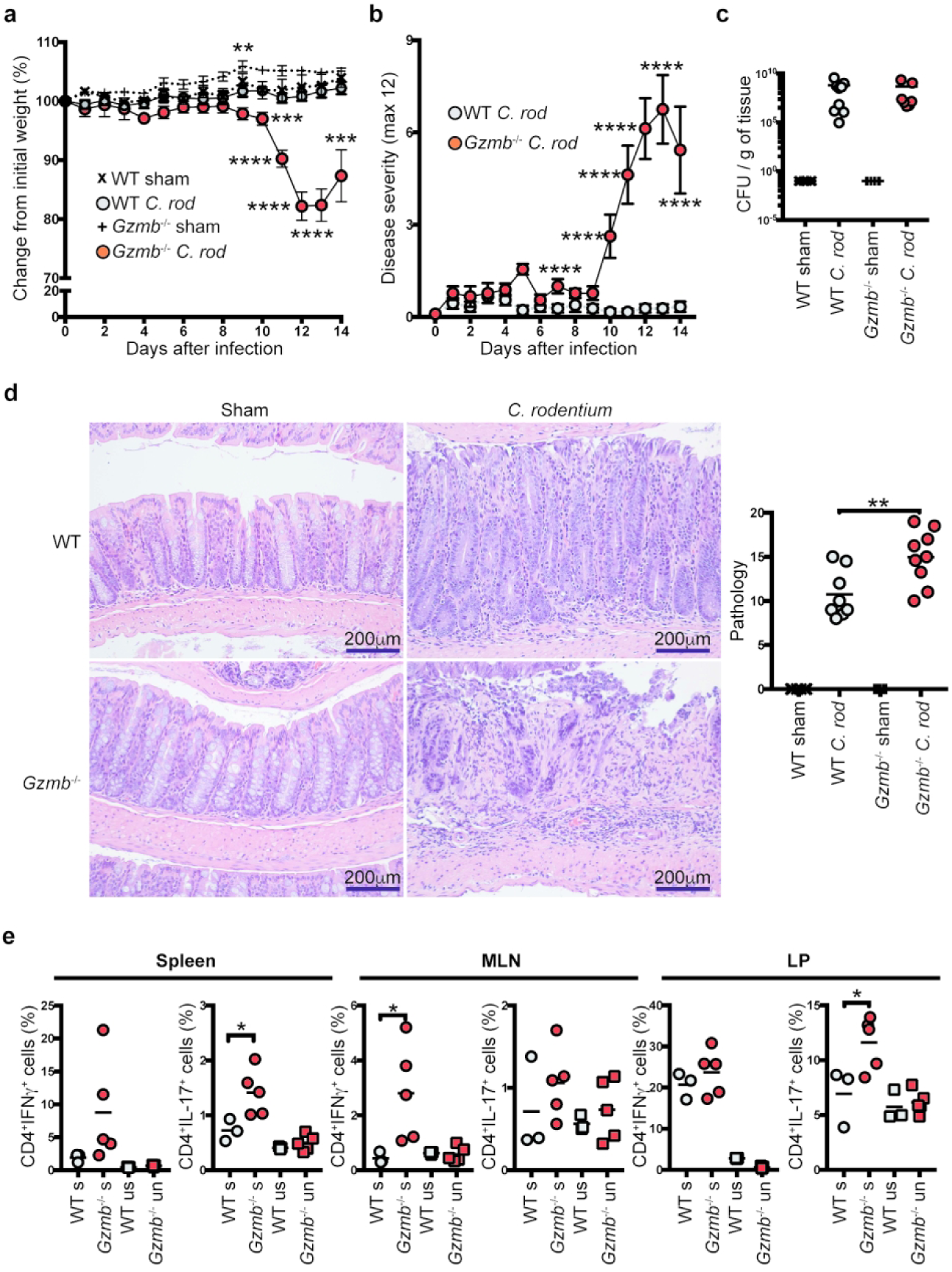
Granzyme B-deficient mice present severe disease after *C. rodentium* infection. Indicated mice were orogastrically infected with *C. rodentium* (5×10^8^ CFU/mouse) and monitored for (a) weight change, and (b) signs of disease. At 14 days post infection, colons were dissected, (c) colonization/g of colon was determined, and (d) pathology scored as indicated in the Methods section; left, representative micrographs (200X magnification); right, data summary. (e) CD4^+^ T cell IFNγ and IL-17 expression in the indicated organs was determined. Cells were non-stimulated (ns) or stimulated (s) for 4 h with PMA/ionomycin. Each dot represents an individual mouse. For (a-d), n=7-9; for (e), 3-5. Data are representative of at least 3 independent experiments. For (a): **P<0.01; ***P<0.01; ****P<0.001; Student’s t-Test. For (b): ****P<0.001; Student’s t-Test. For (d) and (e): *P<0.05; **P<0.01; One-way ANOVA.

Because innate immune cells are important for the response against *C. rodentium*, we investigated whether the absence of granzyme B in innate immune cells was responsible for the observed increased pathology in infected *Gzmb*^-/-^ mice. For this purpose, *Rag-2*^-/-^ and *Rag-2*^-/-^ *Gzmb*^-/-^ mice were infected with *C. rodentium* and monitored for 14 days. Both groups of mice maintained similar weights throughout the course of the experiment (Supplemental Figure 5a), indicating that the effect caused by granzyme B-deficiency is most likely associated with adaptive immune cells. To further confirm these results, we treated *Rag-2*^-/-^ and *Rag-2*^-/-^*Gzmb*^-/-^ mice with anti-CD40 antibodies, which induce acute intestinal inflammation in the absence of T or B cells.^33^ As shown in Supplemental Figure 5b, *Rag-2*^-/-^ and *Rag-2*^-/-^*Gzmb*^-/-^ mice presented similar weight loss throughout the course of the experiment.

Interestingly, when *Gzmb*^+/-^ and *Gzmb*^-/-^ littermate mice were infected with *C. rodentium*, both groups displayed similar weight curves and colon colonization (Supplemental Figure 5c and 5d). However, there was a slight trend for increased IFNγ^+^IL-17^+^ CD4^+^ T cells in the spleen of *Gzmb*^-/-^ mice (Supplemental Figure 5e).

## Discussion

Granzyme B expression in CD4^+^ T cells is well-documented, and has been primarily studied in the context of granzyme B/perforin-dependent regulatory T cell suppressor activity.^34^ However, as our data indicate, granzyme B expression appears to be a common feature among a significant fraction of activated CD4^+^ T cells. Interestingly, not all differentiation conditions induced granzyme B expression. For example, Th0 and Th1 cells rapidly express granzyme B starting at 1-day post activation, whereas Th17 cells do not express this enzyme at any time after activation. In the former conditions, expression of granzyme B is primarily driven by the presence of irradiated splenocytes, indicating the possibility that cell-to-cell contact is important for granzyme B expression.

Although none of the cytokines involved in Th1 differentiation induced granzyme B expression above the levels observed with the addition of irradiated splenocytes, anti-IFNγ and some cytokines present in the Th17 cocktail, such as IL-6 and TGFβ prevented the expression of this enzyme, which suggests that restriction of granzyme B expression is necessary for Th17 polarization. Studies in cytotoxic CD8^+^ T cells have shown that activation of these cells with anti- CD3 in the presence of IL-6, but not in the presence of TGFβ, induces granzyme B expression.^35^ Here we show that in CD4^+^ T cell differentiation, IL-6 does not enhance the expression of granzyme B, indicating that CD8^+^ and CD4^+^ T cells possess different mechanisms for modulating granzyme B production.

Granzyme B expression may not only be limited during the initial CD4^+^ T cell priming and differentiation. For example, TGFβ is known for its role in regulatory T cell differentiation, and some of these cells require granzyme B for their suppressor functions. However, as our results indicate, TGFβ ablates expression of this enzyme, suggesting that regulatory T cells must acquire granzyme B expression during post-priming events. Similarly, a fraction of effector CD4^+^ T cells that migrate into the intestinal intraepithelial lymphocyte compartment acquire granzyme B expression after transcriptional reprogramming.^36^

One of the most intriguing questions is: Why do CD4^+^ T cells express granzyme B during Th0/Th1 activation and differentiation? During *in vitro* Th0/Th1 differentiation, WT-gzmB^pos^ and WT-gzmB^neg^ cells have distinct IFNγ profiles, suggesting that these cells may represent different lineages and that the activity of granzyme B influences their outcome. This is better exemplified when analyzing granzyme B-deficient CD4^+^ T cells, where *in vivo* and *in vitro* activation skew the cells towards an IL-17-producing phenotype, with a distinct gene expression signature. We postulate that granzyme B-deficient IL-17^+^ cells represent a lineage of cells that underwent aberrant differentiation. Therefore, expression of granzyme B during Th0/Th1 activation serves as a checkpoint for proper CD4^+^ T cell differentiation, preventing IL-17 production and increased pathology. These observations have important significance because it has been reported that allelic variants of granzyme B correlate with autoimmune disorders,^37–39^ raising the possibility that in certain CD4^+^ T cell-mediated disorders, lack of proper granzyme B function increases the probability of improper differentiation and increased pathogenesis potential. If this idea is correct, then WT-gzmB^pos^ CD4^+^ T cells may represent cells in which the differentiation signals have the potential to skew cells into an unwanted phenotype, and granzyme B expression in these cells ensures the correct differentiation process.

We have shown that adoptive transfer of CD4^+^ T cells derived from non-littermate (Figure 2) and littermate (Supplemental Figure 2a) mice resulted in increased pathogenicity and IL-17 production, which argues that granzyme B deficiency has an intrinsic effect in CD4^+^ T cells. However, the host’s environment also influences the pathogenicity and IL-17 production of granzyme B-deficient CD4^+^ T cells. For example, infection of *Gzmb*^-/-^ mice with *C. rodentium* resulted in more severe disease with increased IL-17 production relative to WT mice (Figure 6); however, this effect was not observed in infected *Gzmb*^+/-^ and *Gzmb*^-/-^ littermate mice (Supplemental Figure 5c-5e). This observation suggests that granzyme B deficiency in CD4^+^ T cells is necessary but not sufficient for their associated increased IL-17 production and pathogenicity. One potential factor that may be responsible for controlling the pathogenicity of granzyme B-deficient CD4^+^ T cells may be the microbiota present in granzyme B-competent mice. We are currently investigating this interesting hypothesis.

Granzyme B activity in lymphocyte development and/or differentiation has been previously reported.^40^ These authors showed that granzyme A and B are important for the development of a subset of TCR^neg^ intestinal intraepithelial lymphocytes known for the expression of intracellular CD3γ. Development of these cells requires granzyme B to cleave and inactivate the intracellular domain of Notch1. This raises the interesting possibility that during CD4^+^ T cell differentiation, granzyme B prevents unwanted phenotypes by disrupting signals coming from the environment, such those provided by Notch and its ligands, which are known to influence CD4^+^ T cell differentiation.^41^ Our observation that granzyme B expression was increased in the presence of irradiated splenocytes supports this hypothesis. We are currently investigating what signals derived from APC enhance granzyme B expression.

It has been reported that granzyme B-deficient CD4^+^CD25^-^ T cells used in murine models of graft versus host disease (GVHD) induced faster disease onset with increased lethality in comparison to their WT counterparts.^42^ Although this group did not provide a mechanistic explanation for the increased pathogenicity observed, they suggested that increased proliferation and decreased cell death may be involved. According to our data, we believe that in the GVHD model, differential proliferation and survival of granzyme B-deficient CD4^+^ T cells are the result of aberrant CD4^+^ T cell differentiation, which allows escape of highly pathogenic clones. Further investigations in the GVHD model are needed to test this hypothesis.

In summary, granzyme B has been known for its role in cell mediated-cytotoxicity, and while many groups have reported potential extracellular roles for this enzyme,^43^ here we present *in vitro* and *in vivo* evidence supporting a novel intrinsic function for granzyme B during CD4^+^ T cell differentiation. Although many questions remain to be answered regarding how granzyme B is involved in this process, our results provide a strong foundation for a better understanding of the function of this enzyme in CD4^+^ T cell biology.

## Methods

### Mice

C57BL/6J and C57BL/6J.CD45.1 mice were originally purchased from The Jackson Laboratory (000664, and 002014 respectively) and have been maintained and acclimated in our colony for several years. Granzyme B (*Gzmb*)*^-/-^* mice were kindly provided by Dr. Xuefang Cao. Littermate mice were generated by crossing *Gzmb*^-/-^ with WT mice; F1 males (*Gzmb*^+/-^) were subsequently crossed to *Gzmb*^-/-^ female mice to obtain *Gzmb*^+/-^ and *Gzmb*^-/-^ mice. CD57BL/6J *Rag2*^-/-^ mice were kindly provided by Dr. Luc Van Kaer. *Rag2*^-/-^*Gzmb*^-/-^ were generated in our colony. All mice were between 6 to 10 weeks of age at the time of experimentation. Male and female mice were used for all experiments. Mice were maintained in accordance with the Institutional Animal Care and Use Committee at Vanderbilt University.

### Reagents and flow cytometry

Fluorochrome-coupled anti-mouse CD4 (GK1.5), CD8α (53-6.7), CD45RB (C363.16A), granzyme B (NGZB), IFNγ (XMG1.2), IL-17a (TC11-18H10), Ki67 (solA15), TCRβ (H57-597) and ghost viability dyes were purchased from ThermoFisher, BD Bioscences or Tonbo. Biotinylated anti-CD8α (53-1.7) and CD19 (1D3) antibodies were purchased from Tonbo. The Invitrogen Vybrant CFDA SE cell tracer kit was purchased from ThermoFisher Scientific. Anti-CD40 antibody was purchased from BioXcell. Surface cell staining was performed following conventional techniques. For intracellular cytokine staining, cells were stimulated with PMA and ionomycin in the presence of Golgi Stop (BD Biosciences) for 4hr prior to staining. Extracellular markers were stained, cells were fixed briefly with 2% paraformaldehyde, followed by permeabilization and intracellular staining. For intracellular cytokine and granzyme B staining, the BD Cytofix/Cytoperm kit was used according to the manufacturer’s instructions. For intracellular Ki67 staining, the eBioscience transcription factor staining buffer set was used according to the manufacturer’s instructions. All stained samples were acquired using BD FACS Canto II, 3- or 4-Laser Fortessa, or 5-Laser LSR II flow cytometers (BD Bioscences). Data were analyzed using FlowJo software (Tree Star).

### Lymphocyte isolation

Spleen and MLN lymphocytes were isolated by conventional means. IEL and LP cells were isolated by mechanical disruption as previously reported.^44^ Briefly, after flushing the intestinal contents with cold HBSS and removing excess mucus, the intestines were cut into small pieces (∼1 cm long) and shaken for 45 minutes at 37°C in HBSS supplemented with 5% fetal bovine serum and 2 mM EDTA. Supernatants were recovered and cells isolated using a discontinuous 40/70% Percoll (General Electric) gradient. To obtain lamina propria lymphocytes, intestinal tissue was recovered and digested with collagenase (187.5 U/ml, Sigma) and DNase I (0.6 U /ml, Sigma). Cells were isolated using a discontinuous 40/70% Percoll gradient.

### *In vitro* CD4 T cell activation/differentiation

Naïve CD4 T cells were isolated from the spleens of the indicated mice by magnetic sorting using the Miltenyi or StemCell Technologies naïve CD4^+^ T cell isolation kits according to the manufacturer’s instructions. Splenocytes or MLN cells used as APC were incubated for 1hr at 37°C, non-adherent cells were removed, adherent cells were washed and resuspended in complete RPMI, and irradiated with 7 Gy (700 rads). 5×10^5^ CD4^+^ T cells with or without 5×10^5^ irradiated APC were incubated in the presence of plate bound anti-CD3 (5μg/ml) and soluble anti-CD28 (2.5μg/ml), for the indicated times, under the following conditions: Th0 (10ng/mL IL-2), Th1 (10ng/mL IL-2, 20ng/mL IL-12, and 10μg/mL anti-IL-4), Th17 (5ng/mL TGFβ, 20ng/mL IL-6, 10ng/mL IL-1β, 10ng/mL IL-23, 10μg/mL anti-IL-4, and 10μg/mL anti-IFNγ), and Th2 (10ng/mL IL-4, 2μg/mL anti-IFNγ, and 2μg/mL anti IL-12).

### Adoptive transfer of naïve CD4 T cells

Splenocytes from WT or *Gzmb*^-/-^ mice were depleted of CD19^+^ and CD8α^+^ cells using magnetic bead sorting (Miltenyi). Enriched cells were stained with antibodies directed against TCRβ, CD4 and CD45RB, and naïve CD4^+^ T cells were sorted as CD4^+^CD45RB^hi^ using a FACSAria III at the Flow Cytometry Shared Resource at VUMC. 1×10^5^ cells were adoptively transferred into the indicated recipient mice. Starting weight was determined prior to injection. Mice were monitored and weighed weekly. At the end time point, mice were sacrificed and donor-derived cells were isolated from the indicated organs or tissues. Colon histopathology was performed in a blind fashion by a GI pathologist (MBP), following established parameters.^45^ For competitive transfer experiments, 1×10^5^ CD4^+^CD45RB^hi^ cells from both WT-CD45.1 and *Gzmb*^-/-^ CD45.2 mice were adoptively co-transferred into *Rag2*^-/-^ mice and analyzed as indicated above.

### Transcription profile analysis

For gene expression analysis, TCRβ^+^CD4^+^ T cells were purified by flow cytometry from a pool of MLN cells comprised of 1 male and 1 female animal with similar disease progression 21 days after adoptive transfer. RNA was isolated using the RNeasy micro kit (Qiagen). cDNA library preparation and RNAseq was performed by the VANTAGE Core at VUMC on an Illumina NovaSeq 6000 (2 x 150 base pair, paired-end reads). The tool Salmon^46^ was used for quantifying the expression of RNA transcripts. The R project software along with the edgeR method^47^ was used for differential expression analysis. For gene set enrichment analysis (GSEA), RNAseq data was ranked according to the t-test statistic. The gene sets curated (C2), GO (C5), immunological signature collection (C7) and hallmarks of cancer (H) of the Molecular Signatures Database (MSigDB) were used for enrichment analysis. GSEA enrichment plots were generated using the GSEA software^48^ from the Broad Institute with 1000 permutations.

### Real time PCR

For *in vitro* analysis, RNA was extracted from MACS-enriched, differentiated CD4^+^ T cells at day 1 and 3 of culture. For *in vivo* analysis, RNA was isolated from flow cytometry purified donor-derived TCRβ^+^CD4^+^ T cells as described above. cDNA was synthesized using the RT2 First Strand kit (Qiagen). Real time PCR was performed using RT2 SYBR Green Mastermix (Qiagen) on an Applied BioSystems Proflex PCR cycler. All primers were purchased from Qiagen.

### *Citrobacter rodentium* infection

Seven-week old female WT and *Gzmb*^-/-^ or *Gzmb*^+/-^ and *Gzmb*^-/-^ littermate animals were infected with *C. rodentium* as previously described^49^. Briefly, mice were infected with 5×10^8^ CFU exponentially grown bacteria by oral gavage. Starting weight was determined prior to gavage. Mice were monitored and weighed daily for 14d. At the end time point, cells were isolated from the indicated organs or tissues. Cellularity was determined, intracellular cytokine staining was performed, and colon histopathology was performed in a blind fashion by a GI pathologist (MBP) as previously described.^50^

### Anti-CD40 disease induction

*Rag-2*^-/-^ and *Rag-2*^-/-^*Gzmb*^-/-^ mice were injected i.p. with 150μg/mouse of anti-CD40 (BioXcell). Mice were monitored and weighed for daily for 7 days.

## Acknowledgments

This work was supported by NIH grant R01DK111671 (D.O-V.); National Library of Medicine T15 LM00745 (M.J.G.); Vanderbilt Training in Cellular, Biochemical and Molecular Sciences Training Program, T32 GM008554-25 (K.G.M.); Vanderbilt Training in Gastroenterology T32 grant NIH T32 DK007673 (A.N.); and the Digestive Disease Research Center at Vanderbilt University Medical Center, NIH grant P30DK058404 (M.B.P., K.T.W., and D.O-V.). Flow cytometry experiments were performed in the VMC Flow Cytometry Shared Resource, supported by the Vanderbilt Ingram cancer Center (P30 CA068485) and the Vanderbilt Digestive Disease Research Center (DK058404). We acknowledge the Translational Pathology Shared Resource supported by NCI/NIH cancer Center Support Grant 5P30 CA68485-19 and the Mouse Metabolic Phenotyping Center Grant 2 U24 DK059637-16. We also thank the Vanderbilt Technologies for Advanced Genomics (VANTAGE) for their invaluable support.

## Figure legends

**Supplemental Figure 1.**
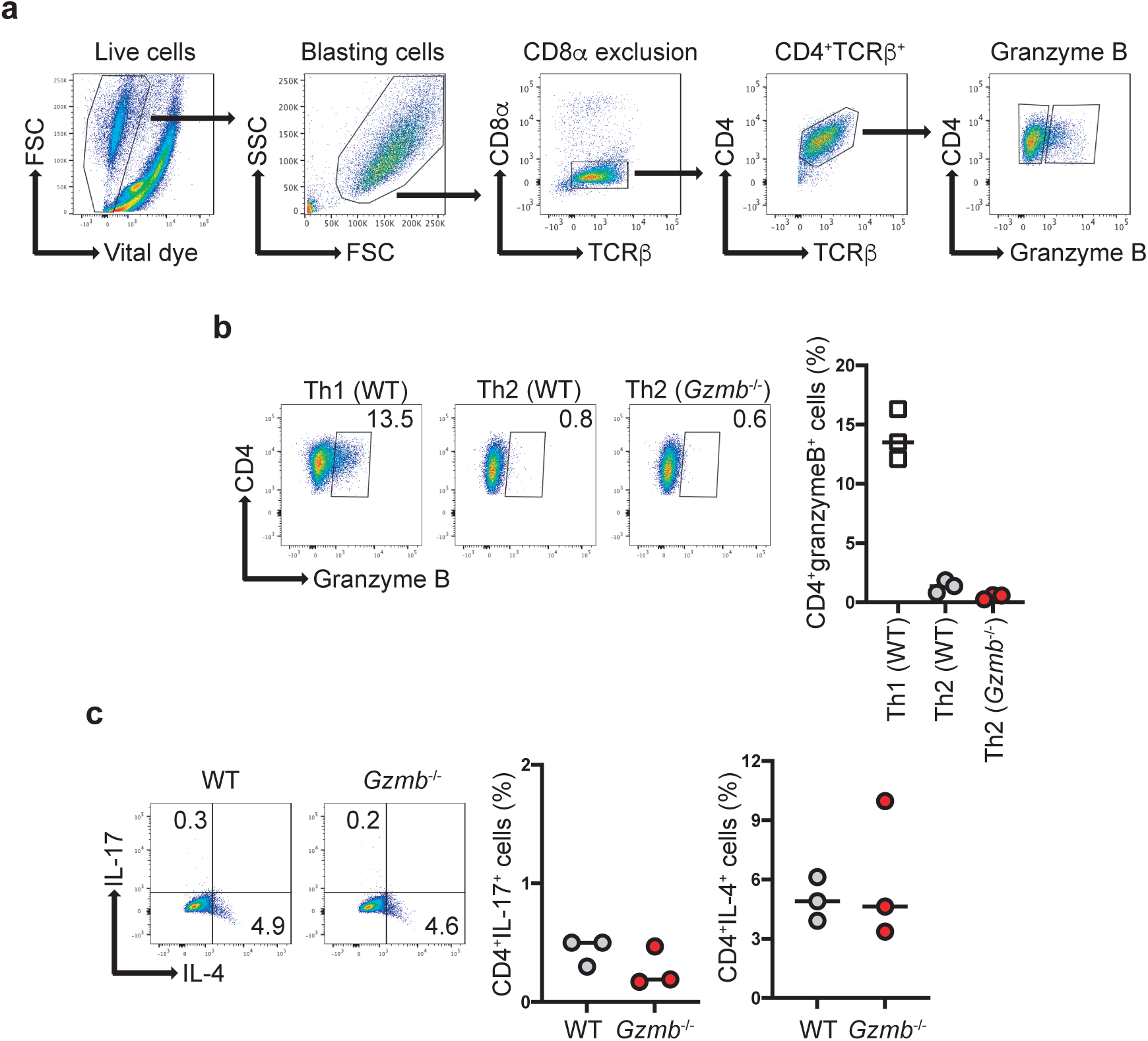
(a) Gating strategy for Figure 1. Plots are from representative WT CD4^+^ T cells cultured in Th0 conditions. Cells from *Gzmb*^-/-^ mice presented similar gating profiles (not shown). (b) Granzyme B expression in differentiated Th1 and Th2 cells from the indicated mice; left, representative dot plots; right, summary. n=3. (c) IL-17 and IL-4 expression in Th2 differentiated cells from the indicated mice; left, representative dot plots; right, summary. n=3.

**Supplemental Figure 2.**
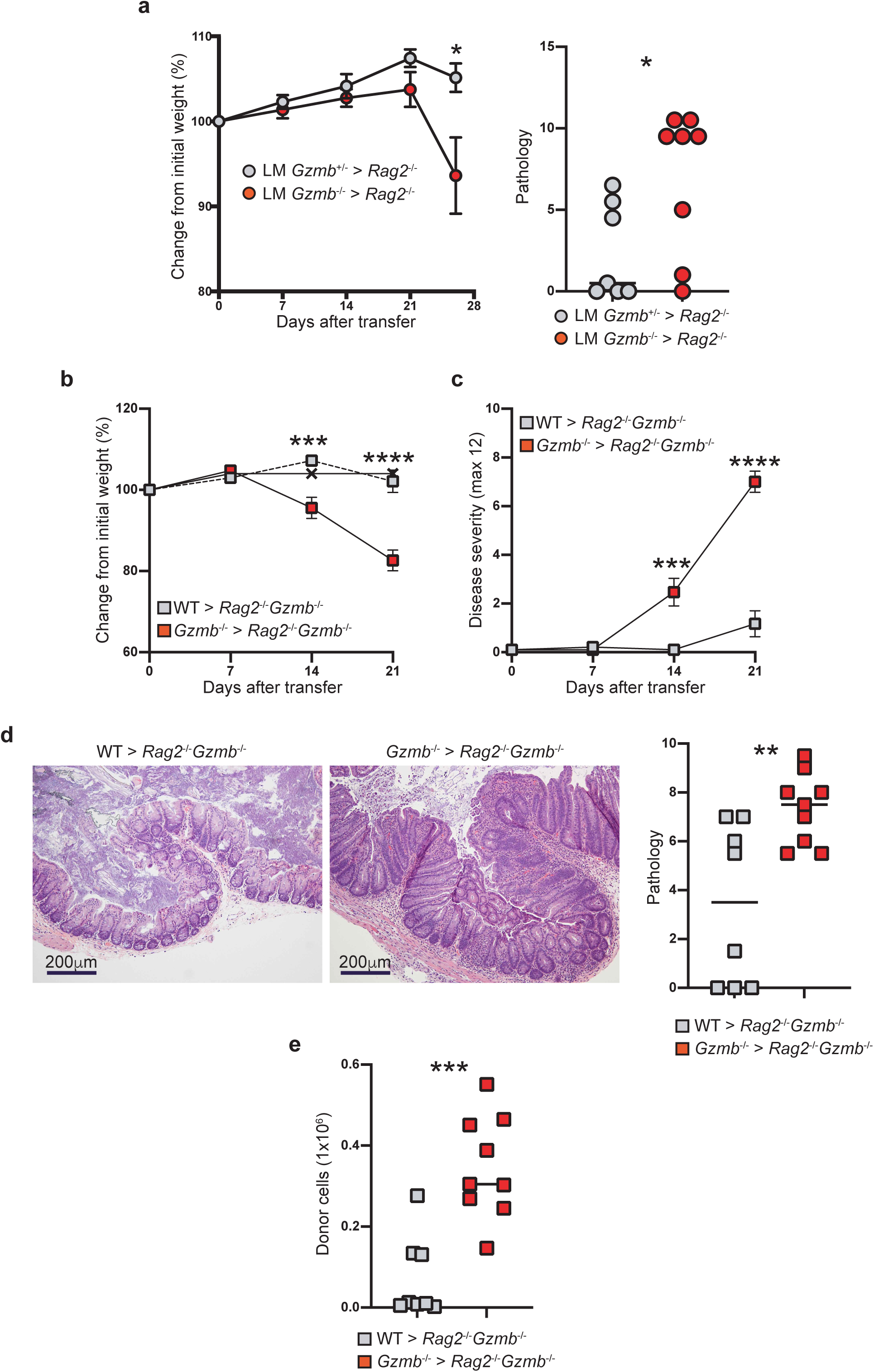
I*n vivo* activated granzyme B-deficient CD4^+^ T cells display increased pathogenicity. (a) 1×10^5^ CD4^+^CD45RB^hi^ T cells from *Gzmb*^+/-^ or *Gzmb*^-/-^ littermate donor mice were transferred into *Rag2*^-/-^ recipient mice and their weights monitored weekly (left), and colon pathology scored (right). (b-e) 1×10^5^ CD4^+^CD45RB^hi^ T cells from WT or *Gzmb*^-/-^ donor mice were adoptively transferred into *Rag2*^-/-^*Gzmb*^-/-^ recipient mice; (b) mice were weighed weekly, and (c) monitored for signs of disease. Twenty-one days after transfer, (d) colon pathology was scored (left, representative micrographs; right, data summary), and (e) donor-cell reconstitution of the IEL compartment in the colon was determined. For (a), dotted lines represent untreated *Rag2*^-/-^*Gzmb*^-/-^ mice. Each dot represents an individual mouse. Data are representative of at least 3 independent experiments. n=8-9. *P<0.05; **P<0.01; ***P<0.001; ****P<0.0001; Student’s t-test.

**Supplemental Figure 3.**
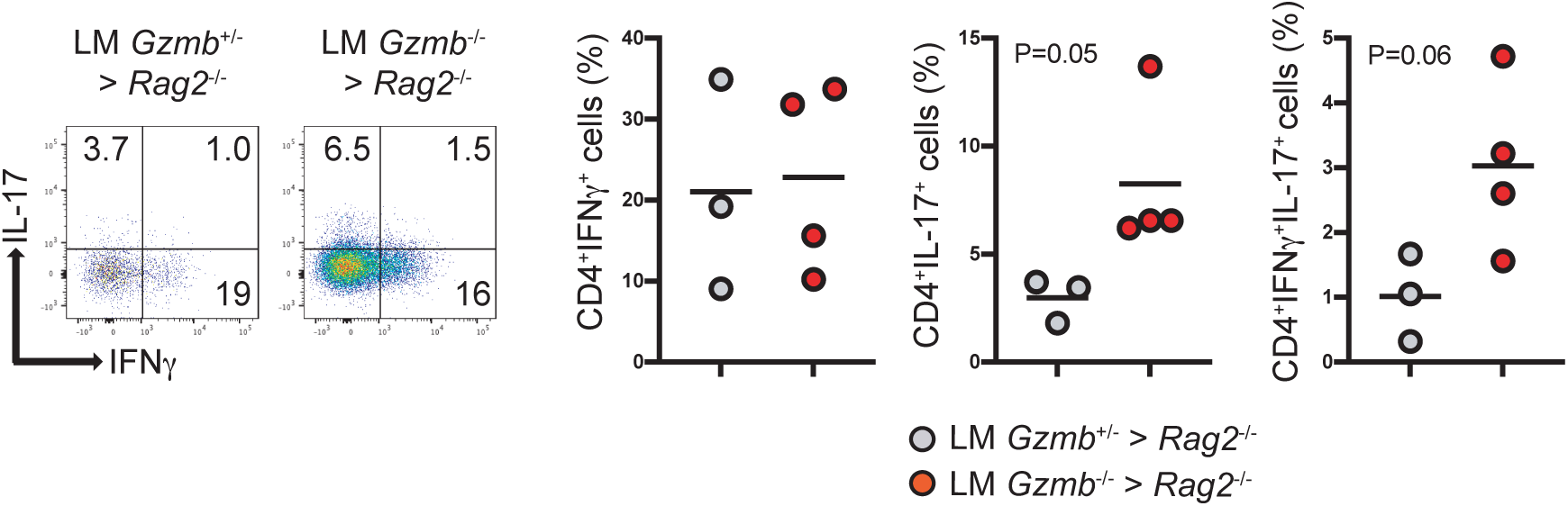
Granzyme B-deficient CD4^+^ T cells present skewed IL-17 differentiation *in vivo.* 1×10^5^ CD4^+^CD45RB^hi^ T cells from littermate *Gzmb*^+/-^ or *Gzmb*^-/-^ donor mice were adoptively transferred into *Rag2*^-/-^ recipient mice. Three weeks after transfer, donor-derived cells were recovered from MLN and their IFNγ/IL-17 profile was determined by intracellular staining. Dot plots indicate a representative sample. Live, TCRβ^+^CD4^+^ cells are displayed. Graphs represent the summary. n=3. Each dot indicates an individual mouse. Student’s t-Test.

**Supplemental Figure 4.**
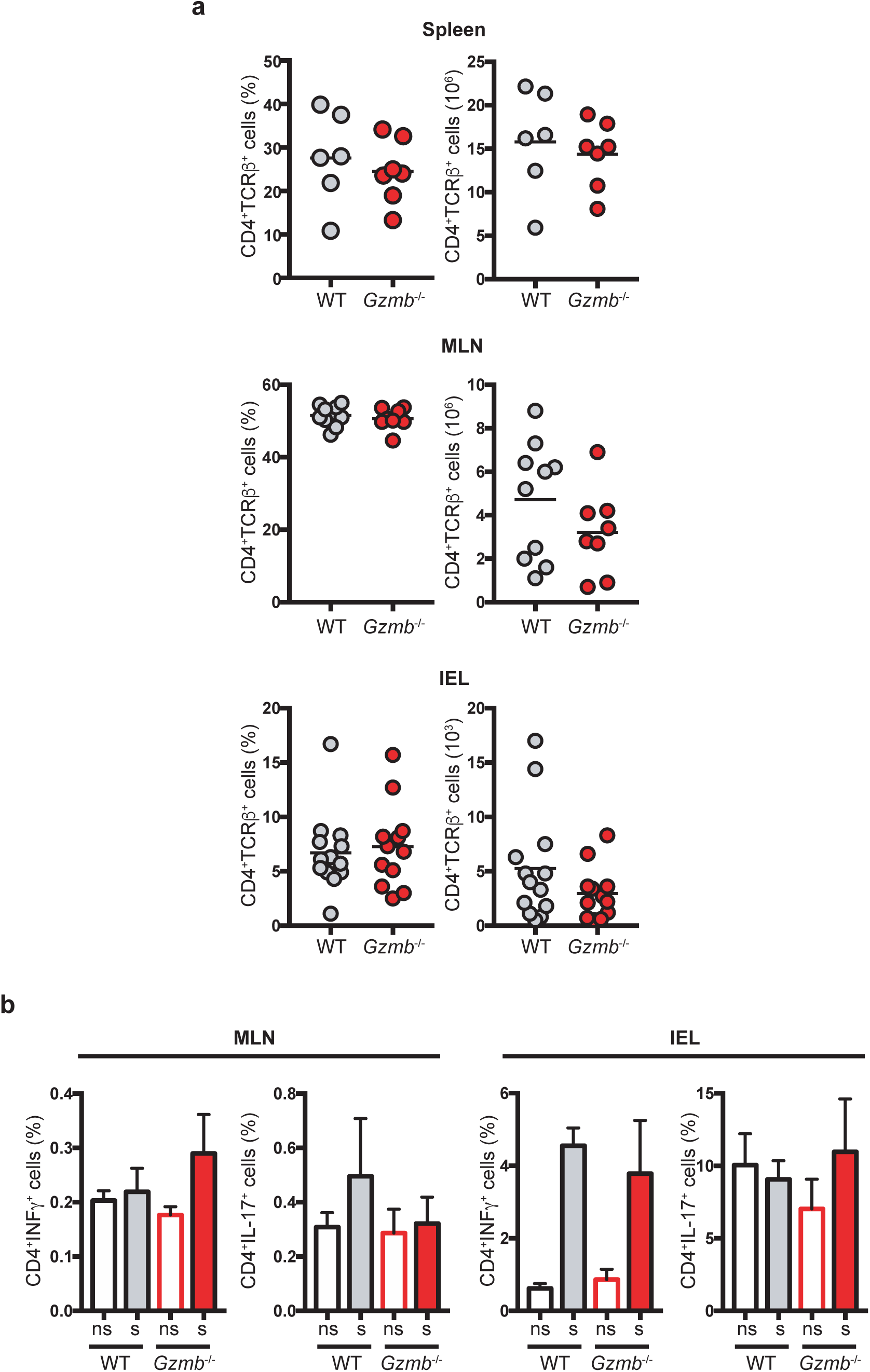
Naïve granzyme B-deficient mice display a normal CD4^+^ T cell compartment. (a) CD4^+^ T cell frequencies and cellularity from spleen, MLN, and IEL compartment were determined by flow cytometry. cells derived from naïve WT and *Gzmb*^-/-^ mice. Cells were non-stimulated (ns) or stimulated (s) for 4 h with PMA/ionomycin. Each dot represents an individual mouse. For (a), spleen and MLN, n=6-10; for IEL, n=12-13. For (b), n=6.

**Supplemental Figure 5.**
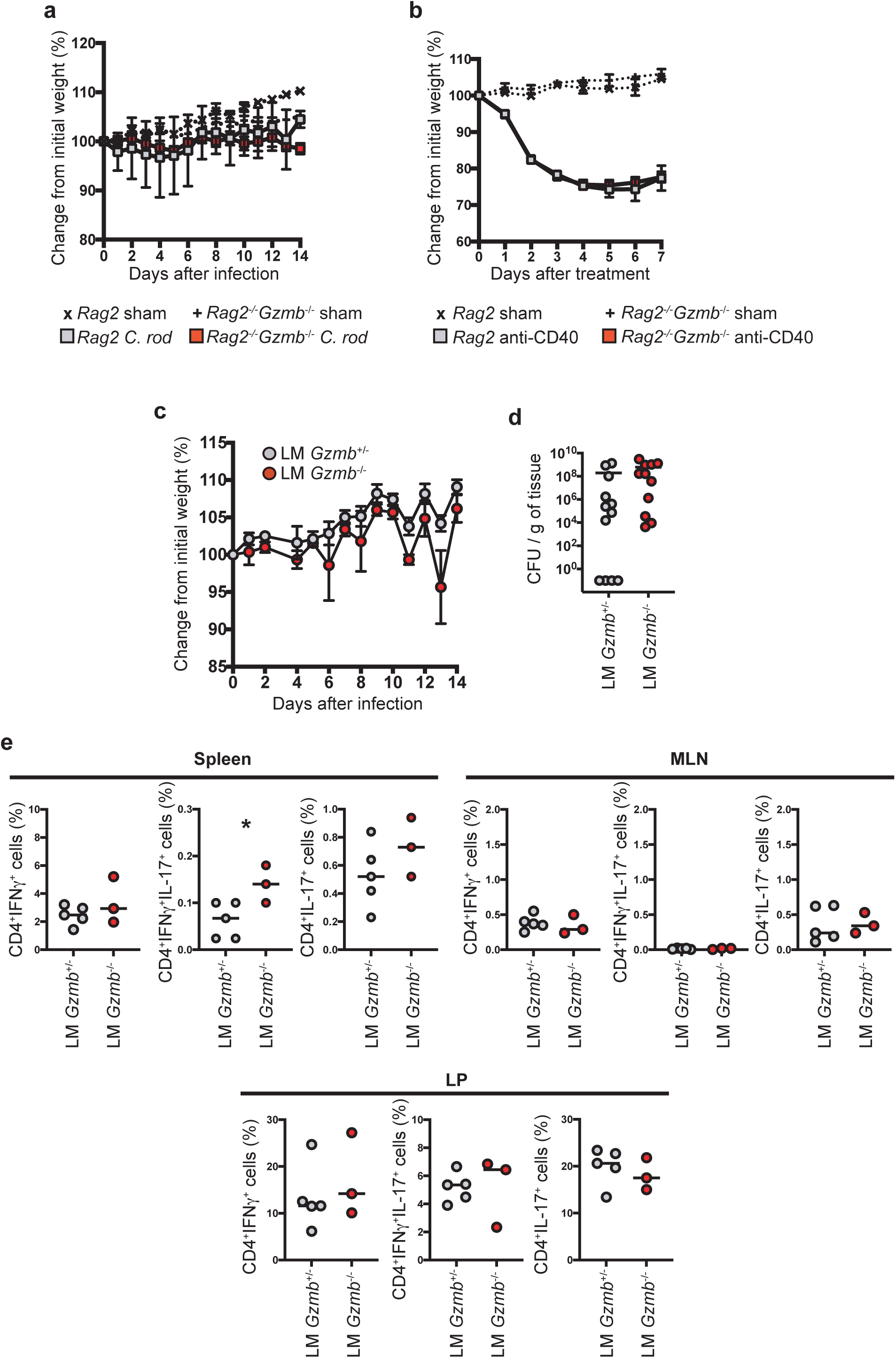
(a) Absence of adaptive immune cells rescues granzyme B-deficient mice from severe disease. The indicated mice were orogastrically infected with *C. rodentium* (5×10^8^ CFU/mouse) and monitored for weight change for 14 days. Data are representative of at least 2 independent experiments. n=3-4 for sham groups. n=5-7 for infected mice. (b) Granzyme B deficiency does not alter disease progression in the anti-CD40 model of colitis. The indicated mice were treated i.p. with 150 μg/mouse and weighed daily. Data are representative of at least 3 independent experiments. n=2 for sham groups. n=14 for treated group. (c-e) Littermate *Gzmb*^+/-^ and *Gzmb*^-/-^ mice were infected with *C. rodentium* (5×10^8^ CF/mouse). (c) Animals were monitored daily for weight change. At 14 days post infection, (d) colon colonization and (e) IFNγ and IL-17 production were determined in the indicated organs. For (c and d), n=11-12; for (e), n=3-5. Student’s t-Test. *P<0.05.

